# Meta-metabolome ecology reveals that geochemistry and microbial functional potential are linked to organic matter development across seven rivers

**DOI:** 10.1101/2024.01.10.575030

**Authors:** Robert E. Danczak, Amy E. Goldman, Mikayla A. Borton, Rosalie K. Chu, Jason G. Toyoda, Vanessa A. Garayburu-Caruso, Emily B. Graham, Joseph W. Morad, Lupita Renteria, Jacqueline R. Hager, Shai Arnon, Scott Brooks, Edo Bar-Zeev, Michael Jones, Nikki Jones, Jorg Lewandowski, Christof Meile, Birgit M. Mueller, John Schalles, Hanna Schulz, Adam Ward, James C. Stegen

## Abstract

Rivers receive substantial dissolved organic matter (DOM) input from the land and transport it to the ocean. As DOM travels through watersheds, it undergoes biotic and abiotic transformations that impact biogeochemical cycles and any subsequent CO_2_ release into the atmosphere. While recent research has increased our mechanistic knowledge of DOM composition within watersheds, DOM development across broad spatial distances and within divergent biomes is under investigated. Here, we combined DOM characterization, geochemical analyses, and shotgun metagenomics to analyze samples from seven rivers ranging from the U.S. Pacific Northwest to Berlin, Germany. Initial analyses revealed that many DOM properties were distinguished by river type (e.g., wastewater, headwater) and that geochemistry often explained variation across rivers. At a global scale, analyses rooted in meta-metabolome ecology indicated that DOM was structured overwhelmingly by deterministic selection. When controlling for scale, however, analyses indicated that ecological assembly dynamics were again partially structured by river type. Finally, microbial analyses revealed that many riverine microbes from our systems shared core metabolic functional potential while differing in peripheral capabilities in across the rivers. Further analysis of the carbon degradation potential for recovered metagenomically assembled genomes indicated that the sampled rivers had strong taxonomically conserved niche differentiation and that carbon degradation potential diversity was significantly related to organic matter diversity. Together, these results help us uncover interconnections between the development of DOM, riverine geochemistry, and microbial functional potential.

## 1. Introduction

Evaluating the principles underlying the composition of river corridor dissolved organic matter (DOM) has been an ongoing, cross-disciplinary endeavor (e.g., ecology, organic matter characterization, biogeochemistry, climatology) (Danczak et al., 2021; Gómez-Gener et al., 2021; Hu et al., 2022; Kaiser et al., 2017; Liu et al., 2019; Xenopoulos et al., 2021). This is because a significant quantity of organic matter is transported through rivers and undergoes combinations of abiotic and biotic transformations that eventually result in the release of CO_2_ into the atmosphere (Cole et al., 2007; Drake et al., 2018; Gómez-Gener et al., 2021; Graham et al., 2018; Regnier et al., 2013; Stegen et al., 2018). By identifying the principles responsible for promoting and constraining DOM composition and incorporating them into various modeling capabilities, we can improve our ability to evaluate watershed function under future conditions (Song et al., 2020; Wohl et al., 2017; Zarnetske et al., 2018).

Microbial communities are one of the more commonly studied factors responsible for influencing DOM dynamics because they can directly interact with various DOM components. Most studies looking at the interconnections between microbial communities and riverine DOM examine a single watershed or reach (Garayburu-Caruso et al., 2020; Graham et al., 2018; Grunert et al., 2021; Osterholz et al., 2016; Saarela et al., 2022; Stegen et al., 2016a, 2018). Within the Columbia River Basin, researchers revealed that mixing dynamics and thermodynamic controls were the key intersections between DOM and microbiology (Graham et al., 2018; Stegen et al., 2016a, 2018). Salinity was an overarching control on both the DOM composition and microbial community structure within the Delaware Estuary, suggesting that external environmental pressures can outweigh the impacts of either assemblage (Osterholz et al., 2016). Studies investigating respiration rates have revealed that DOM thermodynamics play a role in impacting microbial activity, but only when carbon is a limiting factor (Garayburu-Caruso et al., 2020).

Given the heterogeneity of the river corridors above, holistic studies that incorporate information from many different sampling sites are necessary to uncover these principles. Looking at 37 different aquatic environments, Kellerman et al. (2018) revealed that the age of organic matter was consistently related to DOM composition indicating the importance of degradation state. In support of these findings, D’Andrilli et al., (2015) demonstrated that organic matter lability was an important differentiating factor across various aquatic environments, revealing that glacial DOM was the most labile, followed by marine DOM, and finally freshwater. Hu et al. (2022) studied aquatic microcosms across temperature and nutrient gradients and observed that differential assembly processes affected the active and inactive components of DOM, with significant implications for inherent functional traits of DOM molecules. By further expanding both the range of samples and by integrating multiple data streams, we can better uncover important factors impacting DOM composition.

While these studies reveal different factors influencing DOM composition and microbial community structure, these components are often analyzed separately. The recently proposed analytical framework known as ‘meta-metabolome ecology’ represents a potential avenue to develop connections between these data types (Danczak et al., 2020). This analysis treats DOM as an assemblage of molecular formulas subject to processes akin to ecological community assembly processes. By analyzing DOM data this way, researchers can identify transferrable biogeochemical principles and connect biological and chemical data. For example, the use ecological null modeling on DOM helped develop the concept of thermodynamic redundancy, a parallel of functional redundancy observed in microbial communities but for DOM (Danczak et al., 2021). The application of these ecological tools has begun to reveal connections between DOM and microbial communities as well. Similar analyses of both a riverine DOM assemblage and microbial community revealed a lack of selective interconnections, contrary to expectation (Danczak et al., 2020). A combination of DOM null modeling and functional trait analysis indicated that deterministic assembly is predicted to increase under situations of increased global change (e.g., changing temperature/nutrient conditions) (Hu et al., 2022). Turning away from null modeling, functional diversity of DOM assemblages and microbial communities within a freshwater lake revealed significant covariation indicating a potentially underappreciated role in the global carbon cycle (Tanentzap et al., 2019). A study analyzing DOM at a global scale may help reveal transferable principles concerning the behavior of organic matter and is needed because many studies are focused on specific freshwater ecosystems with inconsistent patterns.

Here, we investigated samples from seven rivers spanning a broad geographic extent from the Middle East to the US Pacific Northwest, as part of the Worldwide Hydrobiogeochemical Observation Network for Dynamic Rivers (WHONDRS) consortium. By combining DOM characterization, geochemical analysis, and shotgun metagenomic sequencing partnered to ecological analyses, we assessed environmental interactions which may explain global, river corridor DOM variation and integrate prior observations. In doing so, we sought to characterize the microbes that interact with and alter river corridor DOM. The primary goal of these analyses is to obtain a cross-ecosystem and transferrable understanding of river DOM development and to observe connections between DOM and the microbes that consume it. Ultimately, we observed that the underlying geochemical and lithological conditions appear to be more strongly connected to DOM composition than other factors and that globally distributed microbial species may have conserved niches regarding carbon degradation.

## 2. Materials and Methods

### 2.1 Site Descriptions

This study obtained samples from the seven rivers spanning different biomes and flow regimes (Table 1, Figure 1). These rivers were the Altamaha River (Georgia, USA), Columbia River (Washington, USA), East Fork Poplar Creek (Tennessee, USA), Erpe River (Germany), HJ Andrews – Watershed 1 (Oregon, USA), Jordan River (Israel), and Nisqually River (Washington, USA). The Altamaha River drains a large portion of the state of Georgia to the Atlantic Ocean and has a watershed that extends over approximately 36,000 km2, encompassing the Northern Piedmont and the Coastal Plain (Takagi et al., 2017). The site experiences semidiurnal tidal variations in water levels, with a tidal amplitude on the order of 1-2 m, and occasionally experiences periods of salinity during low flow periods of October. The Columbia River is a gravel bedded river with some riparian vegetation and experiences variable flow associated with dam activity (Arntzen et al., 2006; Stegen et al., 2016a; Villa et al., 2020). The East Fork Poplar Creek has a streambed of gravel, cobble, and bedrock and is surrounded primarily by forest (Demers et al., 2018). This site was historically impacted by mercury contamination because of nuclear weapon development (Brooks and Southworth, 2011; Demers et al., 2018; Loar et al., 2011). The Erpe River is an urban lowland stream that consists of up to 80% treated wastewater originating from an upstream wastewater treatment plant (Mueller et al., 2021). The watershed at HJ Andrews is a shallow, forested, headwater stream that experiences diel fluctuations driven by evapotranspiration (Danczak et al., 2021). The Jordan River is a karstic fed stream found within the northern section of the Syrian African Rift Valley and is subject to discharge fluctuations due to groundwater pumping (Goldman et al., 2019). Finally, the Nisqually River is low-order stream fed by that originates in Mount Rainier and experiences diel fluctuations in discharge due to glacial freeze/thaw cycles (Curran et al., 2016). Additional information and site photos for each river can be found in associated data packages published on ESS-DIVE (see Section 2.8 for complete citations).

**Figure 1:**
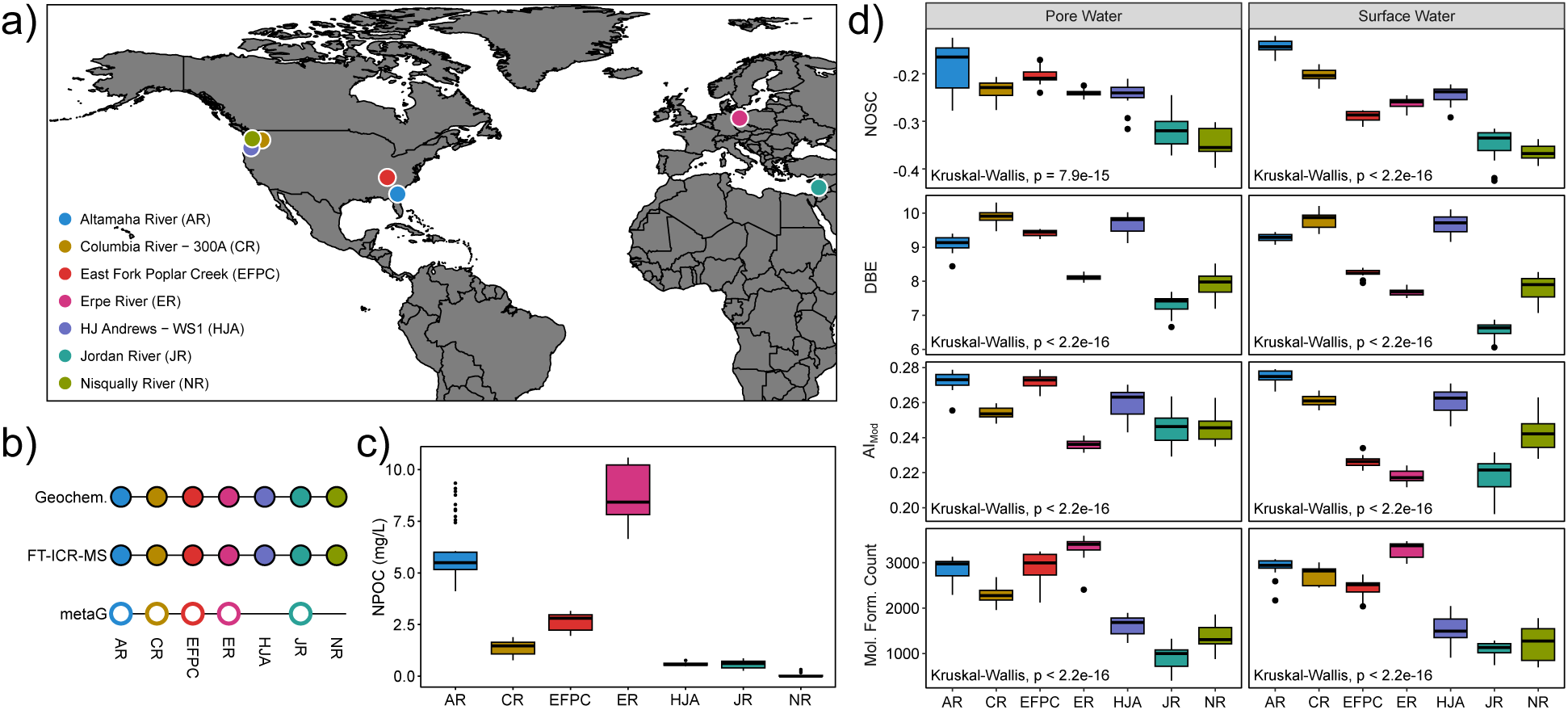
Overview of sampling locations and dissolved organic matter (DOM) characteristics. a) Map of sampling sites. b) Sampling efforts across the seven rivers. Solid circles indicate that all samples were analyzed using the corresponding method, while empty circles indicate a subset of samples was analyzed; no circle indicates no analysis. c) Boxplot of non-purgeable organic carbon (NPOC) concentrations for each river. d) Boxplot of DOM molecular characteristics.

**Table 1:** Summary information for each river. Average pH and non-purgeable organic carbon are provided as contextual information (standard deviations are in parentheticals).

| River | Abbreviation | Location | River Type | pH | NPOC (mg/L) |
| --- | --- | --- | --- | --- | --- |
| Altamaha River | AR | Georgia, United States | Large River | 6 (0) | 5.98 (1.34) |
| Columbia River | CR | Washington, United States | Large River | 7.97 (0.3) | 1.38 (0.32) |
| East Fork Poplar Creek | EFPC | Tennessee, United States | Wastewater | 7.73 (0.37) | 2.60 (0.40) |
| Erpe River | ER | Berlin, Germany | Wastewater | 6.16 (0.37) | 9.5 (1.39) |
| HJ Andrews Watershed 1 | HJA | Oregon, United States | Headwater | 7.79 (0.33) | 0.57 (0.06) |
| Jordan River | JR | Israel | Headwater | 7.72 (0.26) | 0.57 (0.18) |
| Nisqually River | NR | Washington, United States | Headwater | 7.50 (0.25) | 0.06 (0.11) |

### 2.2 Sample Collection and Coordination

Samples from the seven rivers were collected between July and October 2018 as part of a WHONDRS initiative to study diel river dynamics (Stegen and Goldman, 2018). As such, the sampling scheme was similar to the methodology described in Danczak et al., 2021 and was originally devised to investigate temporal patterns despite this study focusing on spatial variation. Three positions separated by ∼4m were selected at each river corridor to collect pore water samples. Approximately 20 mL was collected from each of these locations every 3 h over a 48 h period. Surface water was additionally collected in triplicate from the same position as the middle pore water sampling site. In total, 690 total samples were collected over roughly 17 time points. Surface water was collected using a 60 mL syringe through Teflon tubing while the pore water was collected using a syringe attached via Teflon tubing to a 30 cm long stainless-steel sampling tube (MHE Products, MI, USA) with a slotted screen across the bottom ∼5 cm. One sampling tube was installed to 30 cm depth at each pore water sampling location; these tubes sat in place for around 15 min before the first sample was collected and they remained in place during the 48-h time course of sampling. Prior to sampling a given location, the syringe was flushed 3 times with the source water to ensure only the desired water was collected. All samples were filtered through a 0.2 μm Sterivex filter (Millipore, MA, USA). At each time point, one filter was used for all three pore water samples, and a different filter was used for all three surface water samples. Water passing through the filters was collected for analysis using a needle attached to the filter and injected through a septum. During sampling, water temperature, approximate water stage, and pH were measured. Water samples for DOM analysis were injected into amber borosilicate glass vials. Samples for cations and anions were injected into clear borosilicate glass vials. Once collected, samples were stored in a cooler on blue ice until they could be frozen until they were processed in the lab.

### 2.3 Geochemical Analyses

As with the sampling scheme, the geochemical analyses are similar to the methods described in Danczak et al., 2021. In brief, anion concentrations were measured using a Dionex ICS-2000 ion chromatograph with AS40 autosampler using an isocratic method (guard column: IonPac AG18 guard, 4×50mm; analytical column: IonPac AS18, 4×250mm; suppressor: RFIC ASRS, 300 rmm, self-regenerating; suppressor current: 57 mA). Standards were made from Spex CertiPrep (Metuchen, NJ, 08840) 1000 mg/L anion standards and were diluted as follows: NO_2_ in the range of 0.04 to 20 ppm, F in the range of 0.2 to 10 ppm, Cl and SO_4_ in the range of 0.16 to 80 ppm, and NO_3_ in the range of 0.12 to 60 ppm. Cation samples were prepared with nitric acid and were measured with a Perkin Elmer Optima 2100 DV ICP-OES with an AS93 auto sampler. Calibration standards were made with Ultra Scientific ICP standards (Kingstown, RI): P, Mg, Ca, K, and Na were diluted in the range of 5–4000 ppm, and Fe was diluted in the range of 0.5–400 ppm. Non-purgeable organic carbon (NPOC) was determined by a Shimadzu combustion carbon analyzer TOC-L CSH/CSN E100V with ASI-L auto sampler calibrated to a range of 0.5 to 10 ppm NPOC as C via potassium hydrogen phthalate (Nacalia Tesque, lot M7M4380). An aliquot of sample was acidified with 15% by volume of 2 N ultra-pure HCL and then sparged with carrier gas for 5 min to remove inorganic carbon. The best 3 out of 4 injections replicates were averaged to get the final result.

### 2.4 Dissolved organic matter sample analysis and data processing

All samples were analyzed using a Fourier-Transform Ion Cyclotron Resonance mass spectrometer (FTICR-MS) following methods established previously (Danczak et al., 2021, 2020; Garayburu-Caruso et al., 2020). In brief, samples were standardized to a given carbon concentrations (NPOC 0.69 – 1.5 mg C/L), acidified to pH 2 with 85% phosphoric acid, and extracted with PPL cartridges. High-resolution mass spectra were collected from these samples using a 12 Tesla (12T) Bruker SolariX FTICR mass spectrometer (Bruker, SolariX, Billerica, MA) equipped with an electrospray ionization (ESI) source set to negative mode. BrukerDaltonik Data Analysis (version 5.0) was used to generate a list of m/z values from the raw spectra using the FTMS peak picker module. Chemical formulae were assigned using Formularity (Tolić et al., 2017) according to the Compound Identification Algorithm (Kujawinski and Behn, 2006; Minor et al., 2012; Tfaily et al., 2017) with the following settings: a signal-to-noise ratio >7, mass measurement error <0.5 ppm, and allowing for CHONSP (Formularity formula settings: O > 0 AND (N + S + P) < 6 AND S < 3 AND P < 2).

The R package ftmsRanalysis (Bramer et al., 2020; Bramer and White, 2019) was used to remove peaks that were outside the desired m/z range (200 m/z – 900 m/z) or had an isotopic signature, calculate a number of derived metrics (double-bond equivalent (DBE), modified aromaticity index (AIMod), nominal oxidation state of carbon (NOSC)), and organize the data (Hughey et al., 2001; Koch and Dittmar, 2006; LaRowe and Van Cappellen, 2011; Tfaily et al., 2015). Given the limitations associated with peak intensities, all intensities were converted to presence/absence (Tfaily et al., 2017). Once converted to presence/absence, replicates for each sample were merged such that a peak needed to be present in at least two of the three replicates for it to be considered “present” in that sample. For each sample, we also calculated molecular formula richness as the number of molecular formulas assigned in the respective sample.

### 2.5 Meta-Metabolome Ecology

Following the protocols established in Danczak et al., we generated a transformation-weighted molecular characteristics dendrogram (TWCD) (Danczak et al., 2020). The molecular characteristics dendrogram component of the dendrogram was generated by first measuring the Euclidean distance between molecular formulas based on their derived metrics (e.g., elemental composition, double-bond equivalents, modified aromaticity index, and Kendrick’s defect). These characteristic distances were then weighted by transformation distances associated with the mass of each molecular formula. While described in detail elsewhere (Buser-Young et al., 2023; Danczak et al., 2021, 2020; Fudyma et al., 2021; Graham et al., 2018), the transformation analysis estimates the presence of putative biochemical transformations by measuring the mass differences between identified peaks which are then related to a database of known and commonly observed biochemical transformations. Relationships between peaks are then established by measuring the number of transformations required for one peak to potentially become another. These distances are normalized to a range between 0 and 1 and are used to down weight the molecular characteristic distance. This transformation-weighted molecular characteristic distance matrix is then used as input for an unweighted pair group method with arithmetic mean (UPGMA) hierarchical cluster analysis. The output of this analysis represents total potential relations that could exist across all DOM identified in this study, therefore, this TWCD represents an approximation of global DOM relationships.

The TWCD was used primarily to perform a null modeling analysis known as β-nearest taxon index (βNTI) to estimate the ecological assembly processes impacting the DOM (Danczak et al., 2020; Stegen et al., 2015, 2013, 2012). To perform this analysis, a null distribution of 999 β-mean nearest taxon distance (βMNTD) values were generated and compared to the observed βMNTD value for a given pair of assemblages within each river (e.g., cross-river comparisons were not analyzed). Pairwise comparisons with |βNTI| > 2 indicate that deterministic processes were responsible for observed differences in metabolite composition. In contrast, pairwise comparisons with |βNTI| < 2 indicate that stochastic processes were responsible for observed differences in metabolite composition. Furthermore, the deterministic processes can be separated into two classes. When βNTI > 2, differences in metabolite composition are greater than would be expected by random chance (i.e., greater than the stochastic expectation). This is analogous to ‘variable selection,’ which occurs when deterministic processes drive divergence in composition between a pair of assemblages (Dini-Andreote et al., 2015; Stegen et al., 2015). When βNTI < −2, differences in metabolite composition are less than the stochastic expectation. This is analogous to ‘homogeneous selection,’ which occurs when deterministic processes drive convergence in composition between a pair of assemblages. We would like to highlight that while null modeling- was performed on each river independently, the same dendrogram was used in each analysis and was not pruned. This approach allowed us to take a more global perspective in assembly processes and potentially overcome some of the scale limitations observed in other meta-metabolome ecology studies (Danczak et al., 2023, 2021).

Using a similar approach to the microbial data above, we analyzed alpha diversity and beta diversity using the diversity and vegdist/metaMDS functions, respectively, from the vegan R package (v2.6.4). Additionally, we performed a distance-based redundancy analysis (dbRDA) leveraging Jaccard dissimilarity to evaluate relationships between the meta-metabolome and geochemical variables. First, we limited the effect of co-correlated variables by identifying them using the vif.cca function and manually removing variables with high degrees of co-correlation from our model; variables removed from the model were K, Na, and SO_4_ Next, we performed forward and reverse variable selection to identify the minimum set of geochemical variables that could explain metabolome variation using the ordiR2step function. The final set of selected variables were NPOC, Cl, Mg, TN, NO_3_, F, P, Fe, and pH. Finally, we evaluated the significance of the final dbRDA utilizing a permuted ANOVA and visualized the RDA with loadings.

### 2.6 DNA Extraction and Sequencing

DNA was extracted from filters using Zymobiomic’s Quick-DNA Fecal/Soil Microbe Microprep kit (cat# 6012) using the manufacturer’s protocol and the soil option. Metagenomic libraries were prepared using the Tecan Ovation Ultralow System V2 and were sequenced on the NovaSEQ6000 platform on a S4 flow cell at Genomics Shared Resource, Colorado Cancer Center, Denver, CO, USA.

### 2.7 Metagenome Assembly, Binning, and Annotation

All raw reads were trimmed for length and quality using Sickle v1.33 (Joshi and Fass, 2011). To maximize bin recovery and quality, we used multiple assembly methods which are denoted in each assembly and bin name as follows: (B) megahit v1.1.1, (C) IDBA-UD 1.1.0, (D) metaspades v3.13.0, (E) bbcms followed by megahit v1.1.1, and (F) subsample reads (25%) using bbtools followed by IDBA-UD 1.1.0 (Bushnell, 2020; Li et al., 2015; Nurk et al., 2017; Peng et al., 2012). All assemblies were then individually binned using Metabat2 v2.12.1 with default parameters to obtain microbial metagenome assembled genomes (MAGs) (Kang et al., 2015).

MAG completion was estimated using CheckM v1.1.2 (Parks et al., 2015). To ensure only quality MAGs were utilized for metabolic analyses, we discarded all MAGs that had completion <50% and contamination >10%. This equates to “high quality” (HQ) bins and “medium quality” (MQ) bin cutoffs outlined by the genome consortium standard. The resulting MAGs were manually curated and then dereplicated using dRep with default parameters to result in a final set of 302 MAGs (>99% ANI which represents strain-level MAG distinctions) (Olm et al., 2017) (Supplemental Table 1). To quantify MAG relative abundance in each temporal sample and condition, trimmed metagenomic reads were mapped to the dereplicated MAG set using bbmap (v38.70) at minid=95, and output as sam files which were converted to sorted bam files using samtools (v1.9) (Bushnell, 2020; Danecek et al., 2021). We required reads to map to at least 75% of a MAG in each sample, and second the MAG had to have at least 3x coverage in that sample. To determine MAGs that had reads mapped to at least 75% of the MAG, we used CoverM (v0.3.2) in genome mode to output MAGs that passed this threshold (–min-covered-fraction 75); MAGs were considered present within a sample if they had mapping above this threshold (Woodcroft, n.d.). To obtain reads per base for each MAG, we used CoverM (v0.3.2) in genome mode to output reads_per_base (reads mapped/genome length), and from this calculated MAG coverage as reads_per_base x 151 bp (Woodcroft, n.d.). MAGs were taxonomically classified and using the Genome Taxonomy Database (GTDB) Toolkit v1.3.0 in August 2020 (Chaumeil et al., 2020). MAG scaffolds were annotated using the DRAM v1.0 tool which uses PFAM (v33.1), KEGG (v89.1), dbCAN (v9), MEROPS (v120), and VOGDB databases (https://vogdb.org/) for annotations (Finn et al., 2016; Kanehisa et al., 2016; Rawlings et al., 2018; Yin et al., 2012). A phylogenetic tree visualization was generated by 1) identifying ribosomal protein S3 (rps3) amino acid sequences in each MAG by leveraging a Hidden Markov Model (HMM) from PFAM, 2) aligning rps3 sequences using MUSCLE (Edgar, 2004), 3) trimming the MUSCLE alignment using trimAl (Capella-Gutiérrez et al., 2009), 4) using the alignment to generate a phylogenetic tree using FastTree (Price et al., 2010), and 5) finally visualizing the tree using the ggtree R package (Yu et al., 2017). Additional analyses were performed on all MAGs using the dbCAN standalone pipeline.

Using the MAG coverage data and the CAZyme count information obtained via the dbCAN analyses, we performed alpha diversity analyses in each sample (diversity, vegan package v2.6.4; Oksanen et al., 2019). Beta-diversity was measured using Bray-Curtis dissimilarity (vegdist, vegan package v2.6.4) and was visualized using non-metric multidimensional scaling (NMDS; metaMDS, vegan package v2.6.4). Microbial taxa at the Order-level and CAZyme identity were subsequently fitted to their respective NMDS graphs using the envfit function in the vegan package (v2.6.4) to variables influencing each sample or MAG.

### 2.8 Statistics and Plot Generation

Statistical analysis and plot generation were performed using R v4.3.1 and RStudio (Posit team, 2023; R Core Team, 2023). Plotting relied on a mixture of functions from the tidyverse collection of R packages (v2.0.0), ggthemes (v4.2.4), ggrepel (v0.9.3), and ggpubr (v0.6.0) (Arnold, 2021; Kassambara, 2023; Slowikowski, 2023; Wickham et al., 2019). Kruskal-Wallis tests were used to evaluate variation across multiple groups (e.g., whether differences existed across all rivers) using the kruskal.test function, and Mann Whitney U tests were used to evaluate pairwise variation using the wilcox.test function. Spearman correlations were performed using the rcorr function from the Hmisc package (v5.1.0) (Harrell, 2023).

All code and relevant files can be found on ESS-DIVE at https://data.ess-dive.lbl.gov/view/doi:10.15485/2319037 (Danczak et al., 2024). All raw data used in this study can be found on ESS-DIVE using the following DOIs: https://data.ess-dive.lbl.gov/view/doi:10.15485/1509695, https://data.ess-dive.lbl.gov/view/doi:10.15485/1577266, https://data.ess-dive.lbl.gov/view/doi:10.15485/1577260, https://data.ess-dive.lbl.gov/view/doi:10.15485/1577265, https://data.ess-dive.lbl.gov/view/doi:10.15485/1577278, https://data.ess-dive.lbl.gov/view/doi:10.15485/1577263, https://data.ess-dive.lbl.gov/view/doi:10.15485/1576995 (Chu et al., 2019; Danczak et al., 2019; Garayburu-Caruso et al., 2019; Goldman et al., 2019; Renteria et al., 2019; Stegen et al., 2019; Wells et al., 2019). Metagenomic data has been uploaded to NCBI and can be accessed via the Bioproject # PRJNA946291.

## 3. Results and Discussion

### 3.1 Globally distributed rivers exhibit strong geographic environmental metabolome signatures despite geochemical commonalities in similar surrounding environments

Geochemical analyses revealed substantial environmental variation across the seven globally distributed rivers. Looking first at the geochemical patterns using a principal component analysis (Supplemental Figure 2), we see that each river has characteristic geochemical parameter values. Both surface water and pore water from the Erpe River (ER) are characterized by higher NPOC, K, SO_4_, Na, Cl, and NO_2_ concentrations, while the surface water of East Fork Poplar Creek (EFPC) has higher Ca, Mg, NO_3_, Total N, and P concentrations. Given that both systems are considered either currently or historically wastewater contaminated (Loar et al., 2011; Mueller et al., 2021), we would anticipate elevated levels of many geochemical parameters like NPOC and NO_3_/NO_2_ while other geochemical parameters like Ca and Mg are likely due to the underlying lithology. The Altamaha River (AR) is characterized by elevated iron concentrations likely due to the dominant sediment mineralogy, lower pH, and higher NPOC (Supplemental Figure 1) (Bhatti et al., 2009; Roebuck et al., 2020; Shi et al., 2001). The four remaining rivers (and the pore water from EFPC) are defined by lower concentrations of most parameters, particularly the HJ Andrews Watershed (HJA) and the Nisqually River (NR) which are a headwater stream and a glacial melt run-off stream, respectively (Curran et al., 2016; Danczak et al., 2021). Examining carbon data for combined pore and surface water samples (Figure 1c), we observe significant differences in NPOC concentrations across the seven rivers resulting in the formation of three broad groups: ER and AR constitute a high NPOC group; the Jordan River (JR), HJA, and NR form a low NPOC group; the Columbia River (CR) and EFPC have intermediate concentrations between these two groups.

Shifting focus to average environmental metabolome characteristics, each of the seven rivers has a distinct organic matter mixture. The average nominal oxidation state of carbon (NOSC), a metric which can track the thermodynamic bioavailability of organic matter, indicated no groups formed among the rivers. Nevertheless, there were significant differences among some pairwise comparisons (e.g., NR was significantly lower than AR), though few differences exist between water types (e.g., EFPC surface water was significantly different from EFPC pore water) (Supplemental Table 2, Figure 1d). Average double bond equivalents (DBE) and modified aromaticity index (AI_Mod_) were used to evaluate potential structural patterns of organic matter across rivers and water types. Patterns based upon DBE are more apparent with AR, CR, and HJA exhibiting higher values, ER, JR, and NR exhibiting lower values, each for both water types; DBE for EFPC was different across the pore and surface water. No clear groupings across rivers are apparent in AI_Mod_ in either surface water or pore water samples though EFPC, ER, and JR have significantly different average aromaticity across surface and pore water. Overall, the less urbanized, headwater rivers (e.g., NR, JR) have lower NOSC and DBE overall, while the larger rivers with many tributaries (e.g., AR, CR) have higher NOSC and DBE. The two wastewater and one of the headwater rivers (HJA) have NOSC and DBE values intermediate between the two end-member groups. While similar river types (e.g., headwater, wastewater) group together on some metrics, the specific location of each river plays a substantial role in shaping average characteristics. This pattern is also in keeping with data indicating that the surrounding land (often land coverage) plays a differentiating role in DOM composition (Danczak et al., 2023; Roebuck et al., 2020). Moreover, these patterns are not exclusively a function of river NPOC concentration.

River type (e.g., wastewater vs. headwater) is a significant determinant for molecular formula richness (e.g., the number of molecular formulas). Rivers fell into two groups based on molecular formula richness – AR, CR, EFPC, and ER form a high richness group, while HJA, JR, and NR constitute a low richness group and these groups held across both water types (Figure 1d). The patterns in richness closely mirror the NPOC concentrations, with some disagreement; namely, EFPC and the CR have higher richness relative to their NPOC concentrations. Likely, this is in part driven by wastewater for EFPC and the large number of tributaries that feed into the CR Specifically, wastewater is typically high in diverse types of organic matter while the many tributaries of the CRB bring the potential for varied organic matter (e.g., pyrogenic DOM from wildfires, farmland input, urban, etc.) despite comparatively low overall concentrations (Danczak et al., 2023; Vaughn et al., 2023). The low formula richness group consists of samples that were collected either in or close to the headwaters. This data may suggest that larger rivers under agricultural/urban influence (CR and AR) and wastewater-impacted rivers (EFPC and ER) will have higher richness than less urbanized headwater streams (HJA, JR, and NR). Such a pattern is consistent with observations made within the Upper Mississippi River basin whereby forested rivers had lower molecular formula richness than urban and agricultural rivers (Vaughn et al., 2023). These results, in conjunction with previous studies in diverse basins, indicate that incorporating the surrounding land into predictions regarding DOM development is critical. However, the specific interactions between land type and environmental conditions and subsequent impacts on DOM are not clear.

### 3.2 Ecological null modeling of environmental metabolomes reveals geographically distinct forces drive the development of DOM in rivers

Multivariate analyses revealed that the DOM composition was significantly different across rivers and both water types (Figure 2a). The exceptions to this pattern were HJA and NR which had statistically indistinguishable surface and pore water samples. Both systems have low NPOC values and HJA has a high degree of hydrological connectivity between surface and pore water (Ward et al., 2018; Wondzell, 2006; Wondzell et al., 2007). Importantly, the HJA results differ from our previous study which indicated that the pore water and surface water metabolomes were significantly divergent (Danczak et al., 2021). The contrasting pattern for HJA between the two studies is likely driven by a shift in scale – by providing more chemodiversity from additional rivers in this study, the surface and pore water HJA metabolomes become much more similar in comparison. Similar results are noted in ecological studies and emphasize that scale has important influences over perceived variation in composition (Hernández, 2020; Ladau and Eloe-Fadrosh, 2019; Levin, 1992; Swenson et al., 2006).

**Figure 2:**
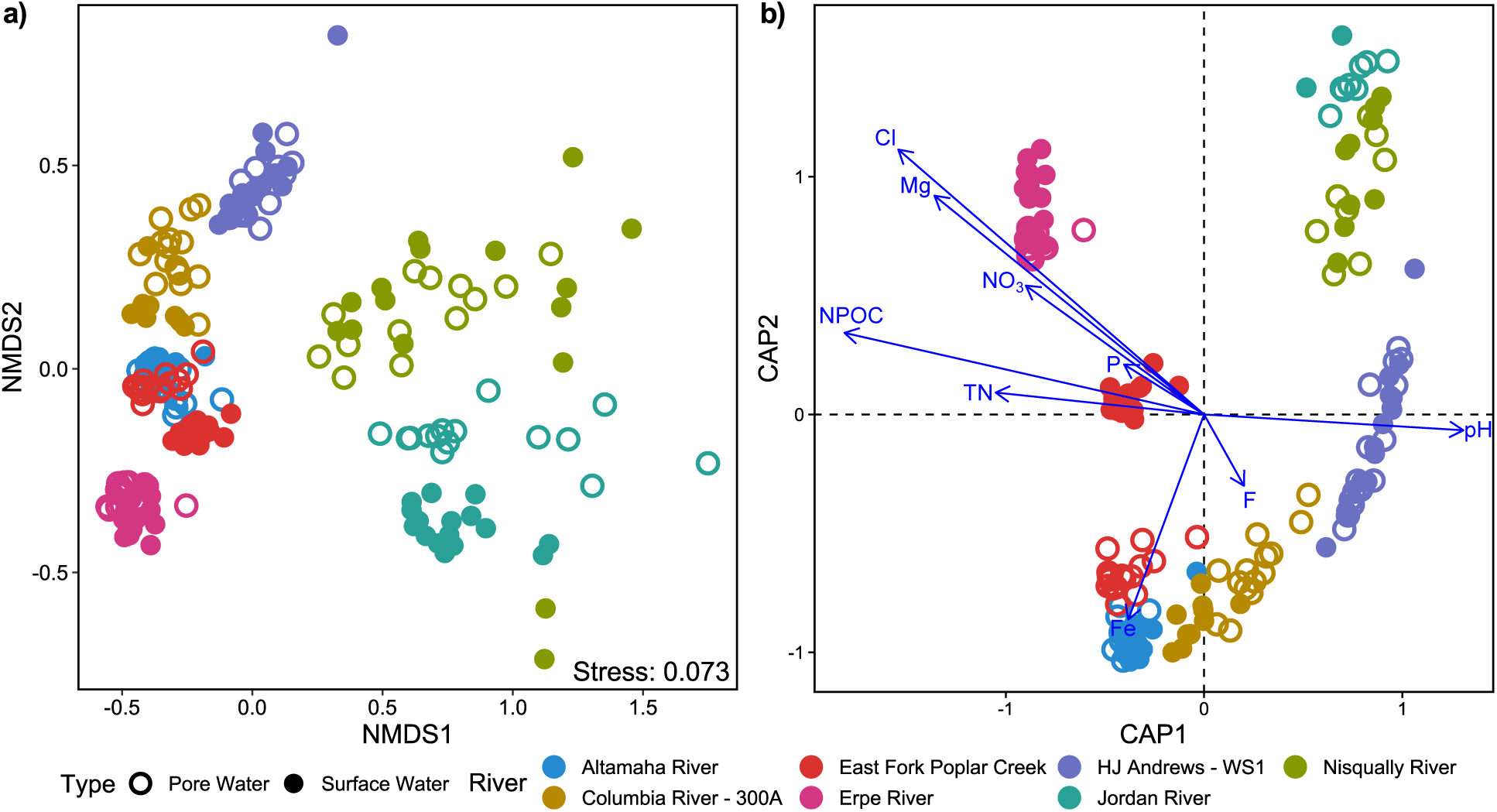
a) Non-metric multidimensional scaling (NMDS) ordination depicting the multivariate relationships of DOM across all seven rivers. b) A redundancy analysis (RDA) optimized to evaluate to what degree the geochemistry is related to DOM composition. Loading arrows indicate geochemical variables significantly related to DOM.

The DOM composition of each river was significantly, but weakly, related to the corresponding geochemical profile (Mantel ρ: 0.1985, p-value: 0.001; PROTEST m_12_^2^: 0.7379, p-value: 0.001). A redundancy analysis (RDA) elucidated which geochemical parameters explained the cross-river metabolome differences (Figure 2b). Initial forward/reverse variable selection prior to the RDA suggested that 9 parameters (NPOC, Cl, Mg, TN, NO_3_, F, P, Fe, and pH) sufficiently captured variability across the samples. The RDA revealed that 6 parameters (Cl, Mg, NO_3_, NPOC, TN, and P) were associated with the samples from wastewater-impacted streams (ER and the surface water of EFPC), while HJA, JR, and NR were associated with higher pH values and lower concentrations of those 6 parameters. As in other analyses, the pore water of EFPC, AR, and CR were intermediate to the other 4 rivers, though there was an iron signal associated with AR and a weak fluoride signal for CR. These relationships indicate a cohesion between geochemical and organic matter variability and suggest that geochemical condition might be a stronger predictor than geographical location (e.g., NR and JR are more similar each other than other geographically closer locations). While useful, these analyses do not tell us how the geochemistry is related to the ecological processes which shape the environmental metabolome.

Following protocols established via meta-metabolome ecology, we observed that rivers experienced significantly different ecological assembly processes (Danczak et al., 2020). Briefly, this method leverages a null model called β-nearest taxon index (βNTI) to identify ecological processes impacting the environmental metabolome. Based upon the βNTI value, we can assign either a deterministic (e.g., selection; |βNTI| > 2) or stochastic (e.g., dispersal; |βNTI| < 2) influence. Deterministic processes can be further separated based upon their sign. When βNTI < - 2, the communities were assembled due to homogenous selection and are more similar than would be expected by random chance. When βNTI > 2, communities experienced variable selection because they are more divergent than would be expected by random chance.

Comparing the environmental metabolomes among rivers, each river experiences variable selection overall (βNTI > 2), with ER having the strongest variable selection and AR having the weakest, bordering on stochastic assembly (Figure 3a). By relating βNTI values to geochemical parameters, we can reveal which parameters are important in structuring environmental metabolomes (Danczak et al., 2021, 2016; Stegen et al., 2016b). This analysis revealed that 8 parameters (Ca, SO_4_, K, Mg, NPOC, Na, NO_2_, and TN in order of correlation strength) were significantly positively related to global βNTI values and iron was negatively related (Supplemental Table 3). Looking to those parameters which were influential in the RDA (Figure 2b), we see overlap between these relationships suggesting that many of the influential parameters may impact ecological processes likely through either metabolic or abiotic interaction. For example, some parameters like calcium, potassium, or sodium might be indicative of underlying lithology (e.g., calcium carbonate content) or anthropogenic influence and suggest that certain geological characteristics might lead to deterministic, variable selection (e.g., larger differences in sodium are related to higher βNTI). Alternatively, other metrics like sulfate, total nitrogen, or nitrite may highlight specific nutrient, metabolic, or respiratory activity related to divergent environmental metabolomes. Iron being the only negative relationship may point to a few possibilities: 1) iron concentrations fluctuate inversely with one of the positively related geochemical variables (e.g., pH), or 2) higher iron concentrations are associated with homogeneous selection in environmental metabolomes via metabolic (e.g., iron reduction) or abiotic (e.g., organometallic) interactions (Lv et al., 2016). Iron’s interpretation is further complicated in that many sites had iron concentrations that were below detection (Supplemental Figure 1). Regardless of the specific interaction, geochemical conditions are significantly related to the ongoing ecological processes shaping environmental metabolomes across seven geographically distinct rivers.

**Figure 3:**
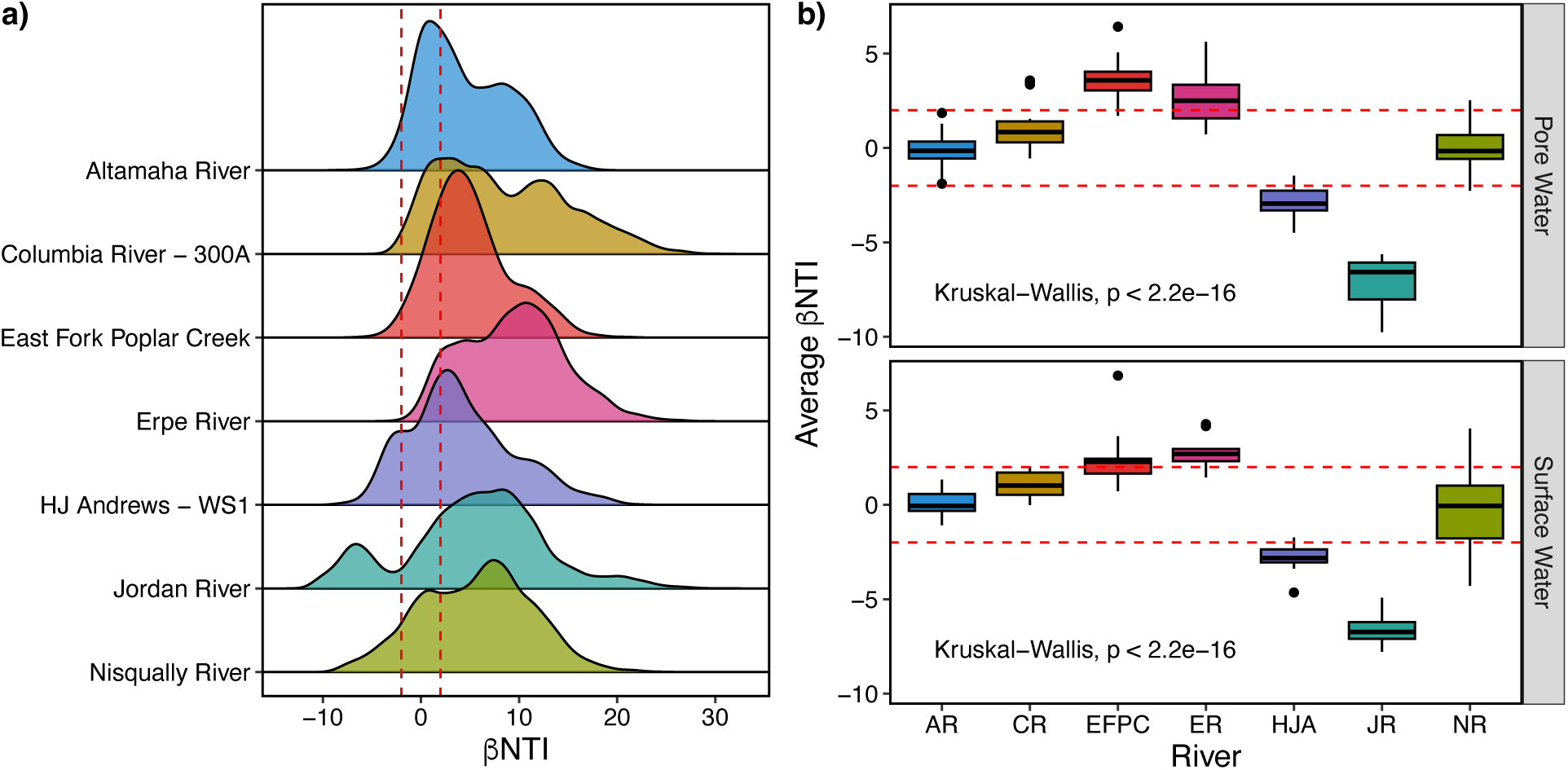
Null modeling results. a) Density plots illustrating the distribution of raw pairwise βNTI values from each River. b) Average βNTI values calculated for pairwise comparisons within the surface water (top panel) or within pore water (bottom panel).

By analyzing βNTI comparisons within a given group (here within rivers), we can evaluate local, rather than global, processes (Figure 3b). EFPC and ER are structured by variable selection (βNTI > 2), HJA and JR are structured by homogeneous selection (βNTI < −2), and AR, CR, and NR are structured by stochastic processes (2 > βNTI > −2). Firstly, we see that the wastewater rivers in this study are governed by distinct ecological assembly processes. Secondly, rivers with low carbon content were more impacted by homogeneous processes. These low carbon rivers also have lower geochemical variability as indicated by principal component and the redundancy analyses. Combined, this suggests that low environmental variability may lead to homogenized organic matter assemblages. Finally, we see stochastic (i.e., dispersal-based) processes impacting rivers regardless of geochemical condition (e.g., low NPOC, low iron NR and high NPOC, high iron AR were impacted by stochastic assembly). These results point to an observation whereby a priori river type appears to be an important characteristic of DOM development in some cases (e.g., ER as wastewater, HJA as headwaters) while dispersal may dominate in others. Similar patterns have been discussed previously whereby land use has been related to DOM diversity (Danczak et al., 2023; Roebuck et al., 2020; Vaughn et al., 2023). Results from these studies support our ecological observations, and indicate that geochemical conditions and the specific location of a river (including the historical influences up to the point in time when sampling occurred) are important in understanding the development of DOM.

### 3.3 The global evolutionary history of river corridor microbes is more significant in dictating carbon degradation potential than local pressures

Microbes are key drivers of biogeochemical cycles and are linked to river corridor organic matter. By understanding the putative functions of resident microbes, we can better understand the forces driving DOM development (Graham et al., 2018, 2016). To evaluate the functional potential as it relates to organic matter diversity, a subset of surface water and pore water samples from AR, CR, EFPC, ER, and JR (1 from each, 10 total) were analyzed using shotgun metagenomic sequencing and a total of 302 metagenomic assembled genomes (MAGs) of medium or higher quality were obtained. These MAGs belong to diverse lineages which are detected across all river corridors (Figure 4a). At higher taxonomic levels, members of the FCB Group and PVC Superphylum were broadly distributed. At lower taxonomic levels, MAGs from the Order Burkholderiales represent at least 1% relative coverage in all rivers aside from JR (Supplemental Figure 3). These Burkholderiales MAGs have consistent carbon, nitrogen, and sulfur processing functional potential, though those found in ER often encoded both an acetate kinase and pyruvate formate lyase and more diverse nitrogen metabolisms (Supplemental Figure 4). This may indicate that these organisms encounter more diverse carbon sources (e.g., short chain fatty acids) and nitrogen types due to their wastewater habitat (Cao et al., 2013; Zhou et al., 2023). While no Burkholderiales were observed within JR, this river shared members of the Order Flavobacteriales with AR, CR, and EFPC and the Order Rhizobiales with AR. The core functional potential of these MAGs is also consistent across the sampled river corridors (Supplemental Figure 4). However, sequences annotated as PEP carboxylase, an enzyme involved in carbon fixation, are present in the Rhizobiales and Flavobacteriales in JR only.

**Figure 4:**
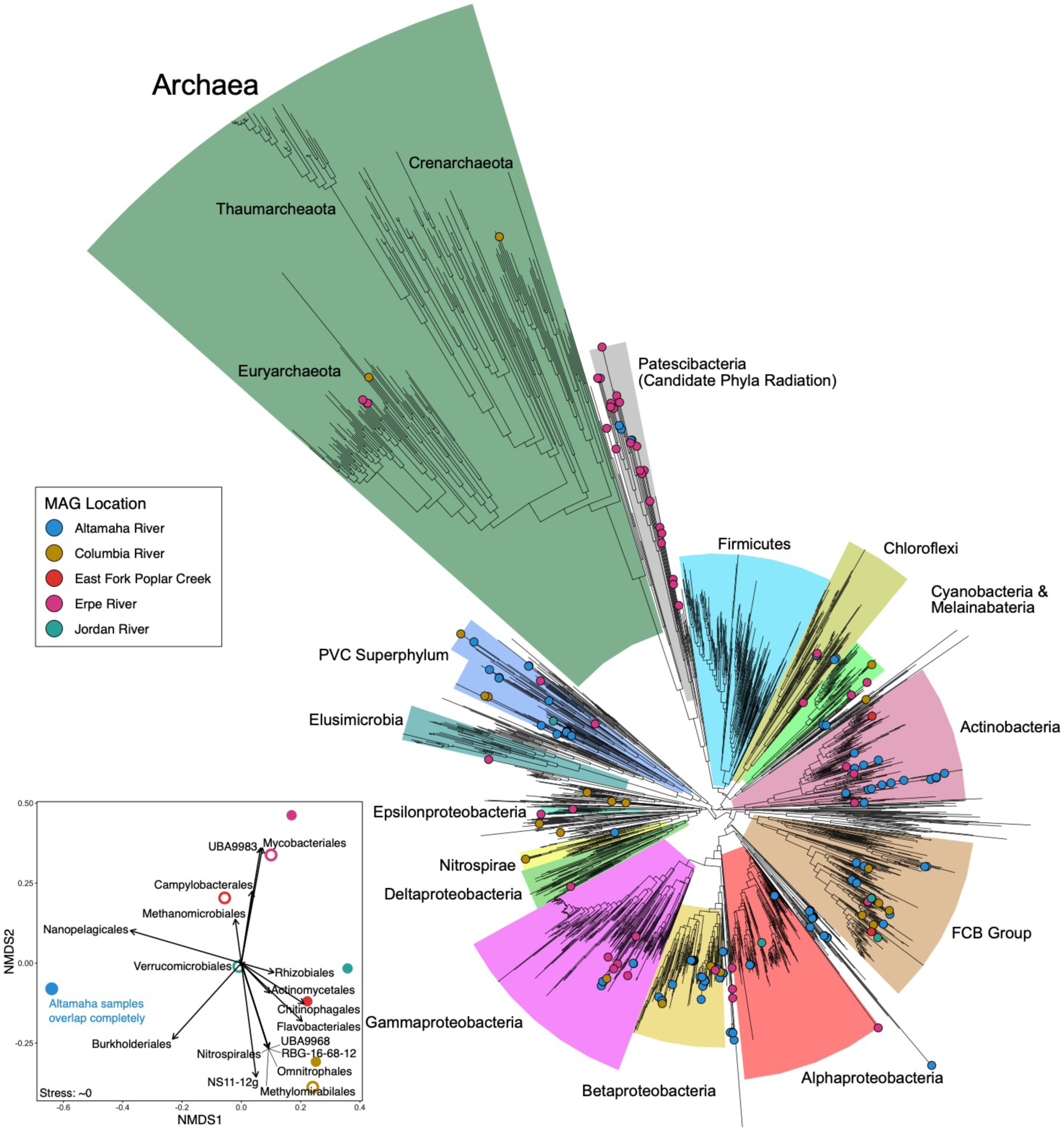
Microbial relationships. a) An rpsC phylogenetic tree generated using FastTree. Points on the tree are genomes obtained from this study and are colored based upon the river from which they were observed. b) Non-metric multidimensional scaling (NMDS) ordination indicating the beta-diversity of each sample. Loadings indicate microbial orders significantly related to the ordination structure.

A similar consistency in core metabolism is observed when examining less broadly distributed taxa (Figure 4b; Supplemental Figure 5). Most Orders (aside from UBA9983) encode common carbon, nitrogen, and sulfur metabolisms with limited variation. The 3 Campylobacterales MAGs found in ER encode pyruvate:ferrodoxin oxidoreductase (PFOR), pyruvate water dikinase, formate dehydrogenase. The Chitinophagales were observed both in CR (4 MAGs) and JR (2 MAGs), but exhibited geographical variations: namely, the JR variants encoded PFOR, the RNF complex, formate-tetrahydrofolate ligase, and a Na-transporting NADH:ubiquinone oxidoreductase. A single Methanomicrobiales MAG was recovered from EFPC and encoded the potential for carbon fixation via the Wood-Ljungdahl Pathway (CO-methylated acetyl-CoA synthase) and for methanogenesis via CO_2_. Many Nanopelagicales MAGs were primarily detected in AR (19 MAGs), though some were observed in CR (1 MAGs) and ER (3 MAGs) with low coverage. These genomes were characterized by the presence of genes encoding an L-lactate dehydrogenase, a partial malonate semialdehyde pathway, a PEP carboxylase, formate-tetrahydrofolate ligase, and 3-phenylpropionate dioxygenase. Finally, the UBA9983 completely lacks genes belonging to the Citric Acid Cycle and has limited metabolic versatility in keeping with other members of the Patescibacteria (also known as the Candidate Phyla Radiation) (Castelle et al., 2017; Vigneron et al., 2023). UBA9983 also exhibits geographical patterns: the 4 MAGs detected in AR encode pyruvate water dikinase and D-lactate dehydrogenases while the 15 ER MAGs encode fewer SCFA enzymes and more diverse nitrogen cycling genes.

These patterns lead to two key observations for these microbial communities: 1) despite significant geographical distances and phylogenetic variation, MAGs observed across the five river corridors have largely similar core metabolic functions, and 2) differences that do arise appear to be related to carbon-type utilization. This type of conservation in functional potential despite phylogenetic variation has been observed in other ecosystems and likely indicates that microbial communities across these five river corridors face many common pressures, though not necessarily dispersal events (Isobe et al., 2020; Martiny et al., 2013; Walkup et al., 2023). In turn, this leads to the development of a common set of functions encoded by varied taxa. However, while many pressures may be similar, each river corridor has a set of unique conditions as evidenced by the geochemical and organic C measurements above (Figure 1; Supplemental Figure 1). This gives rise to a mixture of shared core metabolism and unique peripheral metabolisms, sometimes within the same taxa. For example, while Rhizobiales and Flavobacterales are found in 3 and 4 river corridors, respectively, only those in JR contain a gene characteristic of carbon fixation. The JR also featured some of the lowest NPOC concentrations observed within the dataset. Taken together, this suggests that conditions in the JR might select for carbon fixation due to low carbon concentration. While this is only one example, the other variations in functional potential detected across rivers suggest a more differentiating impact of carbon conditions than other variables (e.g., the differential presence of PFOR, formate-active sequences, etc.).

Diving more deeply into the carbon cycling potential of the Orders described above revealed potential taxonomically conserved niche differentiation across the seven rivers (Figure 5a). Specifically, we see that the Verrucomicrobiales and Rhizobiales (both present in JR) contain more diverse and significantly divergent CAZymes than the UBA9983 (characteristic of ER). An expanded CAZyme analysis for all MAGs revealed that potential niche differentiation exists across the taxonomic groups found in the five river corridors. This is captured by the high cross-taxonomic CAZyme beta-diversity (e.g., distance in the ordination) (Figure 5b). For example, while we see that the Burkholderiales and Nanopelagicales have comparable CAZyme alpha diversity, the distance between them in the ordination is high suggesting divergent roles in environmental carbon degradation. Meanwhile, phylogenetic conservation within taxonomic groups is also suggested by the relatively low within-group beta-diversity. For example, we see that MAGs from the Burkholderiales cluster closely together despite being obtained from 4 geographically distinct rivers (AR, CR, ER, and EFPC). These patterns of taxonomic niche differentiation and phylogenetic conservation of functional potential suggest that the global evolutionary history is more important to an organism’s CAZyme pool than local evolutionary pressures. When combined with the above observation that many differences arise in the functional potential related to smaller carbon compounds, clades are selected for based upon their full carbon degradation potential which can be common across rivers (e.g., Burkholderiales) or specific to an individual river (e.g., UBA9983).

**Figure 5:**
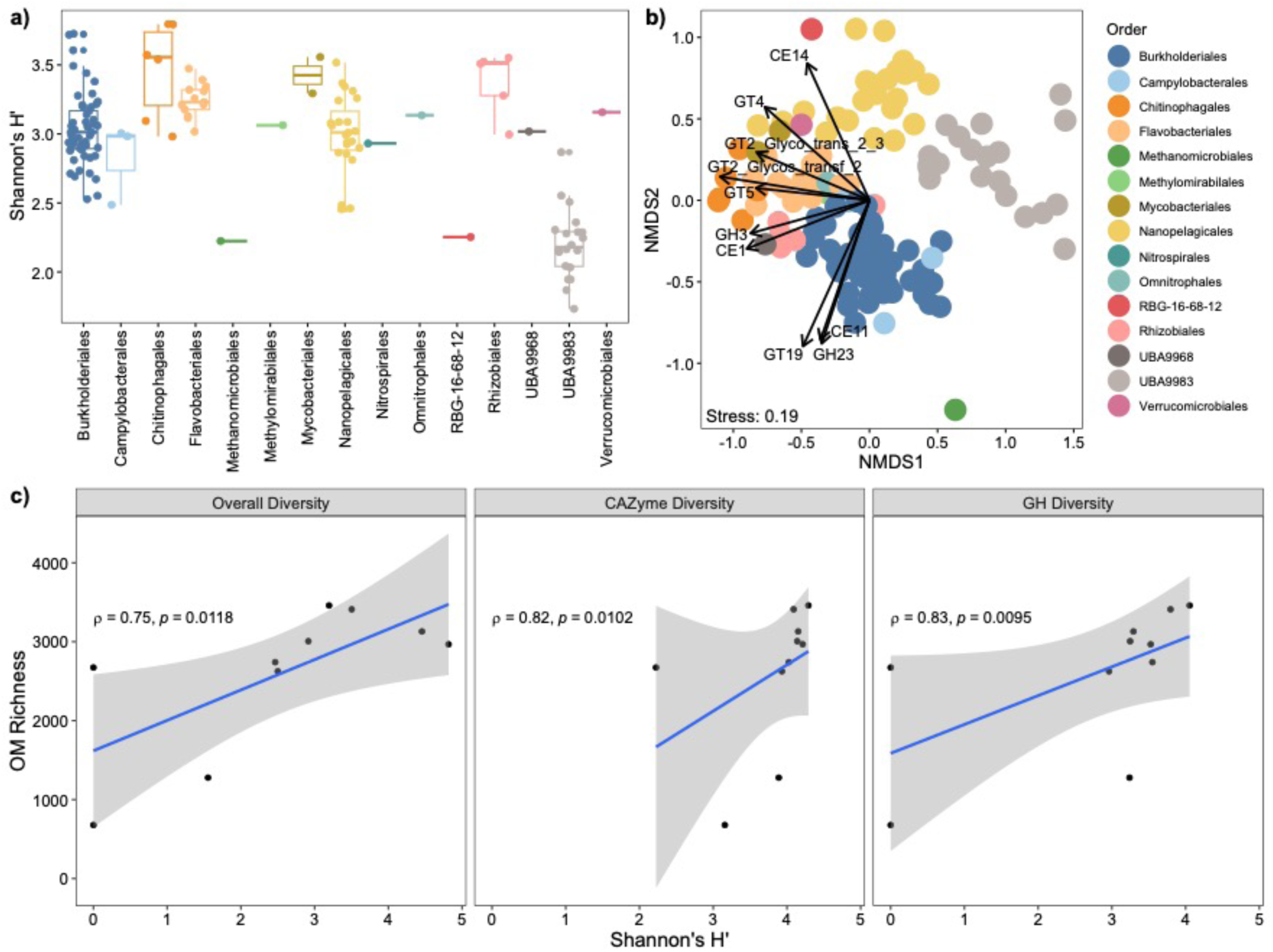
CAZyme analysis. a) Alpha-diversity analysis illustrating the CAZymes pools vary across microbial Orders. b) Non-metric multidimensional scaling (NMDS) ordination of the CAZyme pools illustrating multivariate dynamics across microbial Orders. c) Three different microbially derived diversity metrics plotted against molecular richness.

When various microbial diversity measurements (i.e., Shannon diversity for the overall community, for all CAZYmes contained in the genomes of the community, and for only glycosyl hydrolases in those genomes) are related to DOM richness (i.e., the number of molecular formulas within a sample), we observe significant correlations (Figure 5c). Specifically, the diversity of genes putatively involved in carbon cycling (e.g., CAZymes, glycosyl hydrolases) have slightly stronger relationships with DOM richness than overall diversity (Shannon’s ρ = 0.75 vs. CAZyme and GH ρ = 0.82 – 0.83). This highlights a connection between DOM variety within an ecosystem and the taxonomic and functional potential of microorganisms. When considered in concert with the potential niche partitioning described before, these results suggest that microorganisms may assemble into a community based on the exposure to varied types of DOM, at least in part. Given that carbon cycling potential evolved prior to the dissemination of specific microbial taxa as suggested by the conservation of degradation capability within a given taxonomy (Figure 5ab), DOM might act as a filter for selecting for present groups and may point to causality in the composition of microbial communities. We suggest caution, however, in extrapolating these results too broadly as they were obtained from a limited dataset, from limited data types. Exploration of a greater number of samples across a wider range of biomes while leveraging varied technologies (e.g., metatranscriptomics, metaproteomics, lipidomics, etc.) are necessary before suggesting anything authoritative. Despite these limitations, these results offer a promising glimpse into the ecological interconnections between microbes and the DOM they consume.

## 4. Conclusions

The combination of meta-metabolome ecology, in-depth geochemical analyses, and shotgun metagenomics provided insights into the potential driving forces behind organic matter chemistry and the interconnections between microbial communities and organic matter across seven rivers. The geochemistry of an environment is significantly related to metabolome composition as suggested by traditional multivariate analyses (e.g., Mantel and permuted PROCRUSTES tests) and null modeling approaches. This pattern was expected due to the distinct geochemical signatures of each location. However, our null modeling revealed that there are certain key geochemical parameters that may play a role in structuring organic matter. Calcium, NPOC, sulfate, potassium, sodium, magnesium, total nitrogen, and nitrate consistently were detected as important predictors for variable selection (e.g., positively related to βNTI) (Supplemental Table 4). These geochemical characteristics largely reflect watershed characteristics (e.g. soil characteristics, land cover) and geological processes (e.g., underlying lithology). For example, calcium concentrations in circumneutral conditions are largely controlled by natural weathering processes (Weyhenmeyer et al., 2019). Relationships with calcium (or other primarily weathering derived characteristics) might provide insight into organic matter dynamics that are associated with certain geological histories. They might also point to specific land-use or metabolic processes driving organic matter compositions. Land-use and/or land type (i.e., forested, urban, wastewater, etc.) significantly impacts carbon flux (Regnier et al., 2013) and potentially explains relationships with NPOC. Many of these variables, however, are also involved in numerous metabolic pathways and could point to organismal activity. For example, previous research in HJA and CR revealed that nitrogen metabolism is an important component of DOM processing within those rivers (Danczak et al., 2021; Graham et al., 2018; Stegen et al., 2018). Therefore, the relationships between total nitrogen or nitrate and DOM may represent dynamics in associated nutrient or energy pathways.

Relationships within the microbial communities were more challenging to identify due to the limited sample number, but a few clear patterns emerged. First, we observed members of the Burkholderiales in 4 of the 5 analysed rivers (AR, CR, EFPC, and ER) and members of the Flavobacteriales in 3 of the 5 rivers (AR, CR, and JR). Focusing on carbon cycling potential, it appears that the Burkholderiales play similar roles in each river despite significant geographical distances (Supplementary Figure 3). Second, those lineages which are only found in fewer rivers appear to play diversified roles, potentially related to environmental conditions of a given area. For example, members of the Patescibacteria (UBA9983) are significantly functionally divergent from the Verrucomicrobiales and Rhizobiales. Whereas the Verrucomicrobiales and Rhizobiales appear to have similar carbon cycling potentials as many aerobic organisms, the UBA9983 MAGs encode no respiratory capabilities, have reduced CAZyme diversity, and seem to be obligate fermenters consistent with their taxonomic assignment (Castelle et al., 2017; Vigneron et al., 2023). These differences in putative niche space represent the potential interactions between the microbial community and DOM. While members with a broader potential niche space, like the Rhizobiales or Flavobacteriales, may be able to access more DOM types due to a greater diversity of encoded CAZymes, members with narrower potential niche space (e.g., UBA9983) would be able to target less DOM. When DOM-based diversity measurements are related to these microbial data, we observed significant interconnections highlighting putative role for DOM in selecting for subsets of microbial taxa.

By examining organic matter chemistry in conjunction with geochemistry and microbial functional potential, we were able to identify general trends across seven globally distributed rivers. Namely, we observed that both geochemical conditions and the specific location of a river (i.e., the type of the river, the land use history of the surrounding area, etc.) are important in understanding riverine DOM composition. Additionally, we observed that these same factors significantly influenced the underlying ecological assembly processes structuring DOM, often in a scale-dependent manner.

Finally, our metagenomic analyses indicated that members within the microbial communities are differentiated by their carbon degradation potential with molecular diversity serving as a putative selective filter. Together, these results give us a glimpse into the interconnections between DOM development and microbial functional potential while also highlighting environmental parameters that might be broadly important in understanding DOM in rivers.

## Acknowledgments

Many people were involved in the collection of samples for this project, including but not limited to Xiaojia He, Jurjen Rooze, and Katherine Znotinas. Pacific Northwest National Laboratory is operated by Battelle Memorial Institute for the U.S. Department of Energy under Contract No. DE-AC05-76RL01830. This research was supported by the U.S. Department of Energy (DOE), Office of Biological and Environmental Research (BER), as part of BER’s Subsurface Biogeochemistry Research Program (SBR). This contribution originates from the SBR Scientific Focus Area (SFA) at the Pacific Northwest National Laboratory (PNNL). A portion of this work was performed at the Environmental Molecular Science Laboratory User Facility.

## Author Contributions

R.E.D., A.E.G., and J.C.S. conceptualized the study. V.A.G., J.W.M., L.R., J.R.H, M.J., N.J., B.M.M., H.S., J.S., S.A., E.B-Z. collected samples. V.A.G., J.W.M., L.R., J.R.H analyzed anions/cations. R.K.C and J.G.T. collected FTICR-MS data and assisted with analyses. M.A.B. performed bioinformatic analyses. R.E.D. aggregated data, performed ecological and statistical analyses, and analyzed microbial functional potential. R.E.D. drafted the manuscript but all authors contributed to the writing.

## Supplemental Information

**Supplemental Figure 1:**
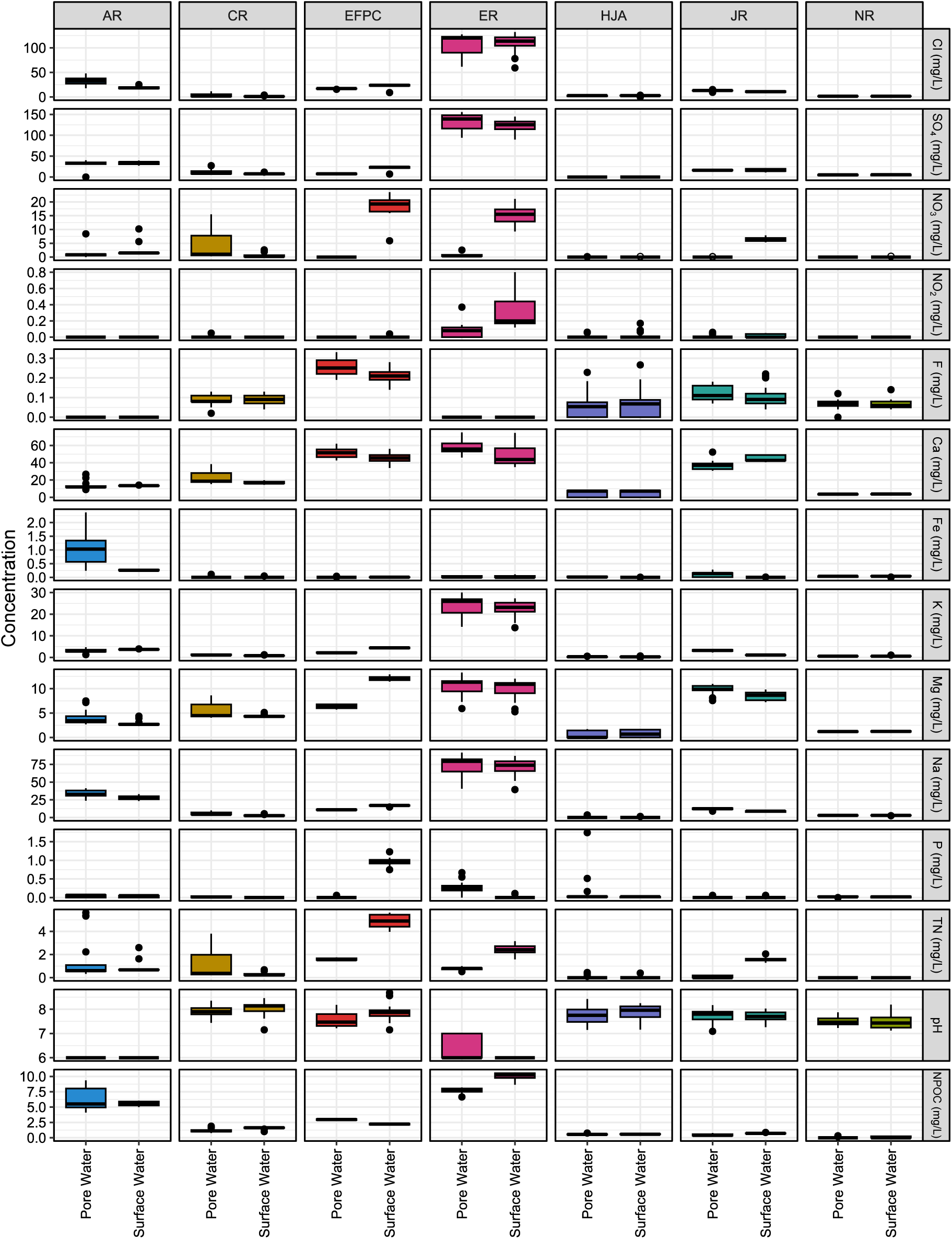
Boxplots displaying variation in geochemical parameters in each river, separated by river type. River abbreviations are AR = Altamaha River, CR = Columbia River, EFPC = East Fork Poplar Creek, ER = Erpe River, HJA = HJ Andrews Watershed 1, JR = Jordan River, and NR = Nisqually River.

**Supplemental Figure 2:**
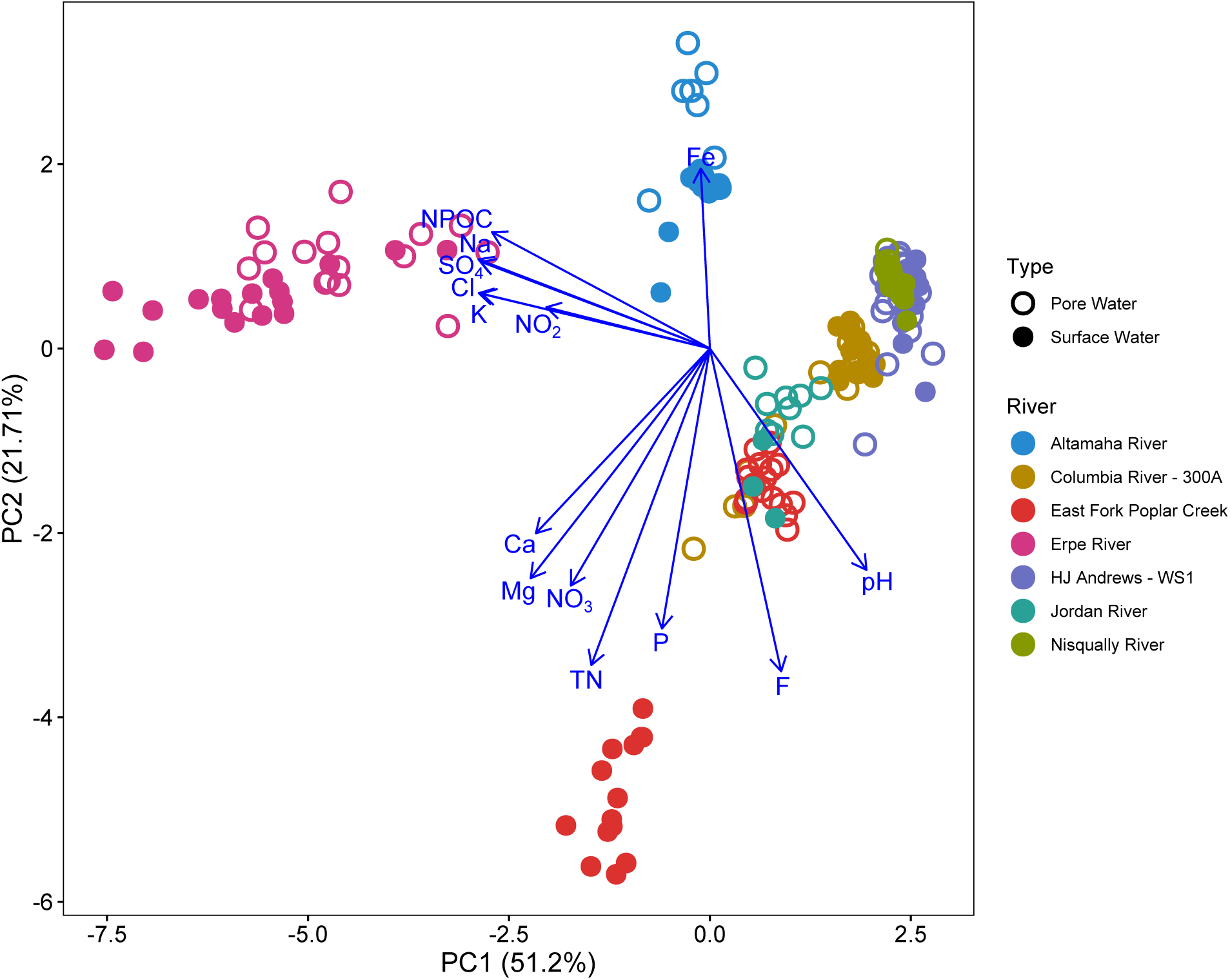
Principal component analysis (PCA) of river and pore water geochemistry.

**Supplemental Figure 3:**
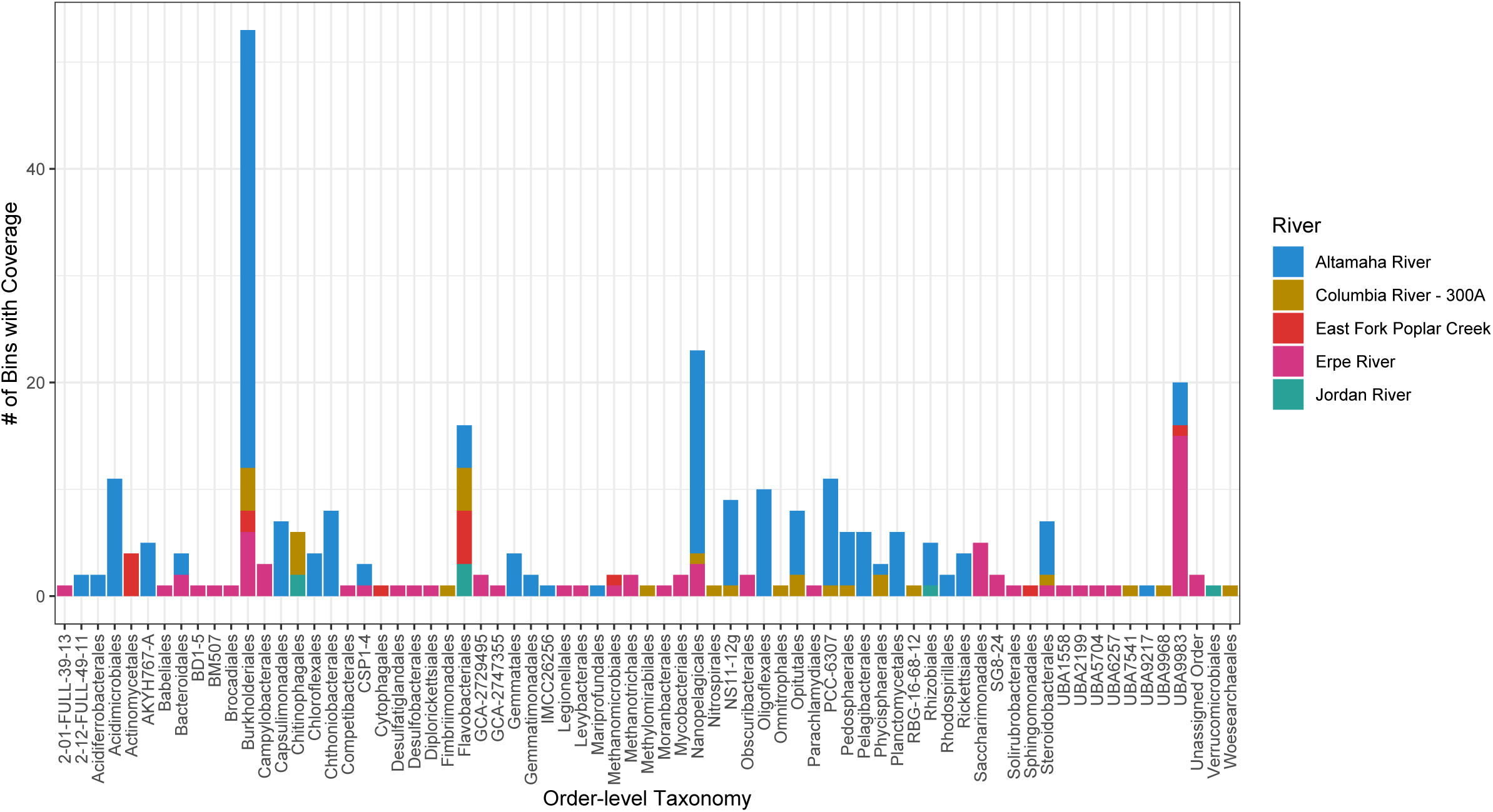
Bar plot depicting the number of bins with a given GTDB taxonomy detected in each river.

**Supplemental Figure 4:**
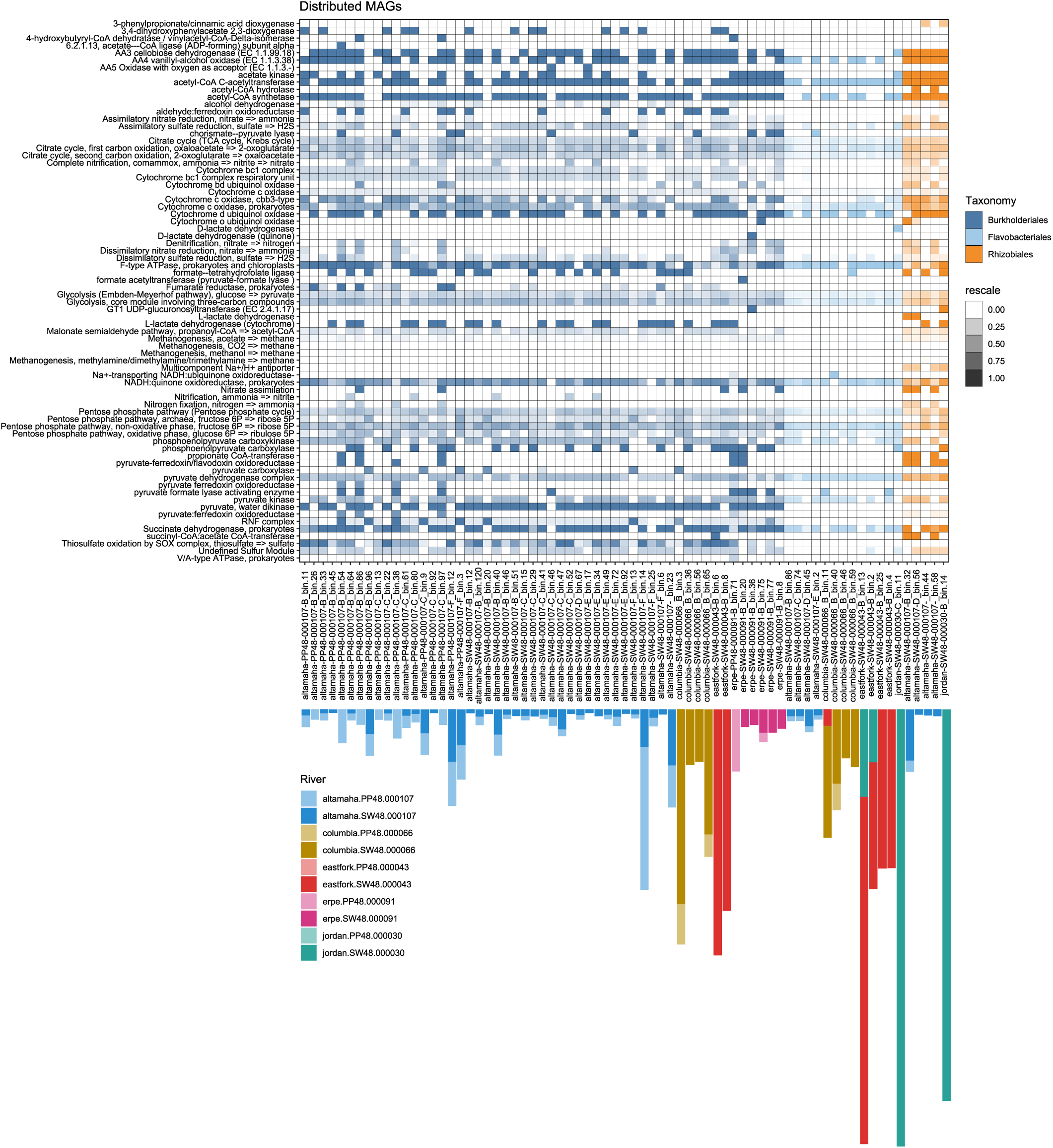
Metabolic information and abundance of broadly distributed MAGs. A heatmap showing the presence (and relative completion) of the pathway listed on the y-axis in the metagenomic assembled genome (MAG) on the x-axis. Cells in the heatmap are colored based upon the taxonomy of the MAG listed on the x-axis. Below each MAG is a visual representation of the read mapping abundance in each river with rivers indicated by the colored bars.

**Supplemental Figure 5:**
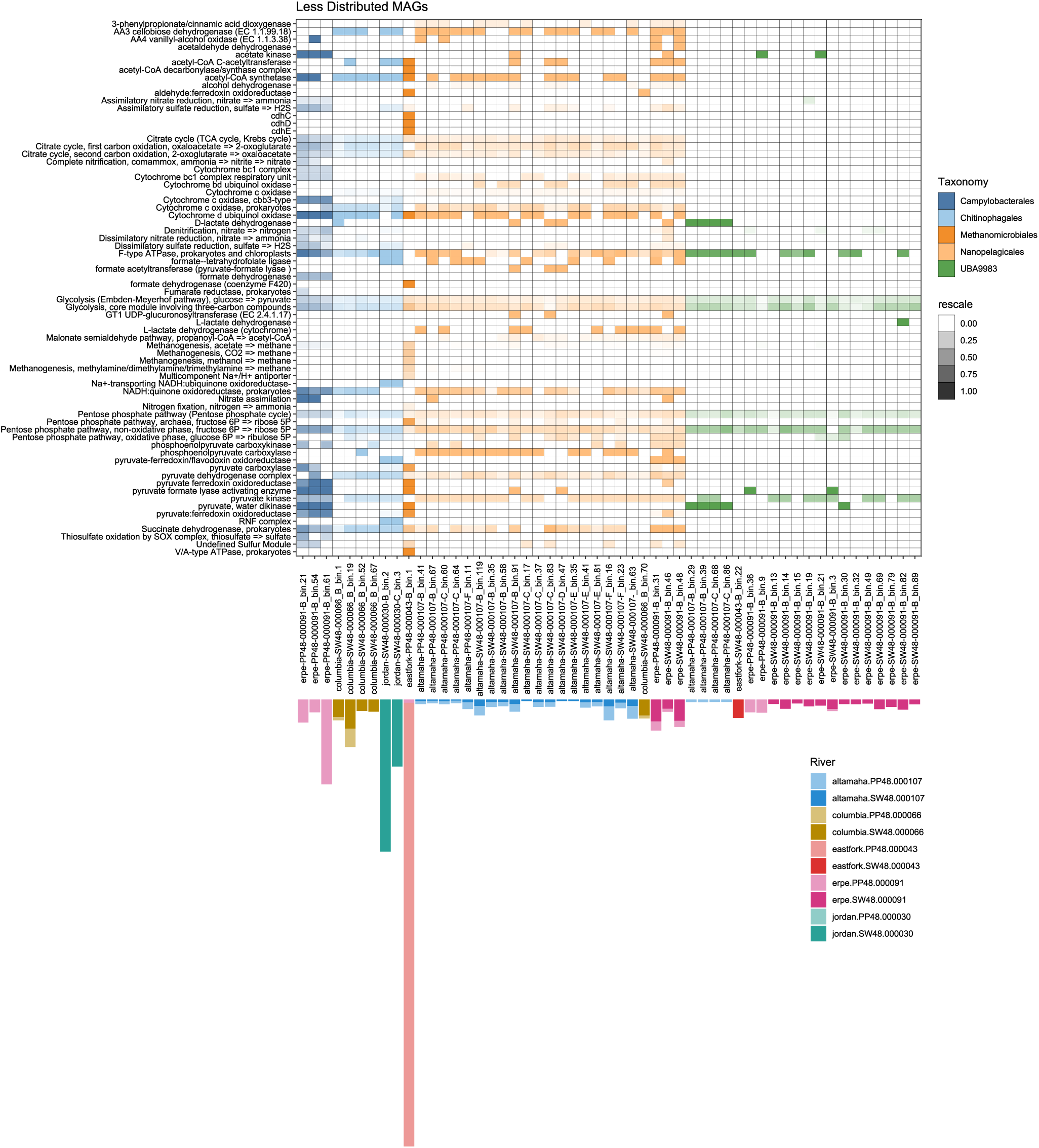
Metabolic information and abundance of MAGs found in fewer rivers (the inverse of Supplemental Figure 4). A heatmap showing the presence (and relative completion) of the pathway listed on the y-axis in the metagenomic assembled genome (MAG) on the x-axis. Cells in the heatmap are colored based upon the taxonomy of the MAG listed on the x-axis. Below each MAG is a visual representation of the read mapping abundance in each river with rivers indicated by the colored bars.

**Supplemental Table 1:** Table describing the summary statistics for each metagenomic assembled genome, including (but not limited to) measurements like completion/contamination percentage, MAG size in base pairs, and N50.

**Supplemental Table 2:** Pairwise Mann Whitney U statistics for DOM measurements across rivers and sample type.

**Supplemental Table 3:** Table of significant correlation values for βNTI and geochemical relationships.

**Supplemental Table 4:** Table of pairwise Mann Whitney U statistics for average βNTI values across rivers.

